# Foreign Ribosome Inactivating Proteins as immune effectors in insects

**DOI:** 10.1101/2023.02.09.527842

**Authors:** Walter J. Lapadula, Maximiliano Juri Ayub

**Affiliations:** Instituto Multidisciplinario de Investigaciones Biológicas de San Luis, IMIBIO-SL-CONICET and Facultad de Química, Bioquímica y Farmacia, Universidad Nacional de San Luis, Ejército de Los Andes, 950, D5700HHW San Luis, Argentina

**Keywords:** Ribosome Inactivating Proteins, Horizontal Gene Transfer, Insects, Immune effectors, RNA *N*-glycosidase

## Abstract

Ribosome inactivating proteins (RIPs) are RNA *N*-glycosidases that depurinate an adenine residue in the conserved alpha-sarcin/ricin loop (SRL) of rRNA. This ribosomal modification inhibits protein synthesis. During the last years, we have reported the existence of these toxins in insects, where their presence is restricted to mosquitoes from the Culicinae subfamily (*e*.*g. Aedes aegypti*) and whiteflies from Aleyrodidae family (*e*.*g. Bemisia tabaci*). Combination of phylogeny and synteny analyses showed that both groups of genes are derived from two independent horizontal gene transfer (HGT) events. Interestingly, we found that RIP encoding genes have been evolving under purifying selection, indicating that they have a positive impact on fitness of host organisms. We also demonstrated that *A. aegypti* RIP genes are transcribed and their transcripts are polyadenylated. Although the biological roles of these toxins remain open to speculation, defense activities have been postulated for plant and bacterial RIPs. Based on these pieces of evidence, we hypothesize that RIPs play a similar protective role in insects. In this work, we report the occurrence of a third HGT event in Sciaroidea superfamily, supporting that RIP genes fulfill an important functional niche in insects. Analysis on transcriptomic experiments from the three groups of insects indicate a convergence in expression profiles which are compatible with immune effectors. Finally, we show the induction in RIP expression after infection with pathogens. Moreover, we show transcriptomic evidence of parasite SRL depurination. Altogether, our results strongly support the role of these foreign genes as immune effectors that confer fitness advantage to host insects.

## Introduction

Horizontal gene transfer (HGT) consists of the non-genealogical transmission of genetic material (Goldenfeld and Woese 2007). The relevance of this mechanism for evolutionary innovation in bacteria is widely accepted (Goldenfeld and Woese 2007; Lerminiaux and Cameron 2019). In contrast, its impact on the fitness of multicellular organisms (*e*.*g*.,animals) is still under debate (Dunning Hotopp et al. 2007; Van Etten and Bhattacharya 2020). For the transferred genes to be permanently maintained in animal species, they must be incorporated into germline cells and transmitted to the offspring. Once vertically transmitted, at least two possible fates are expected for HGT-derived genes. They may be eroded by genetic drift or acquire a functional role in the host species (Keeling and Palmer 2008). While multiple examples of putatively functional HGT-derived genes can be found, little is known about their biological role, how they are regulated in the recipient organism, and ultimately what impact they have on host fitness/survival. In insects like aphids, psyllids, whiteflies, and mealybugs, it has been postulated that HGT has played a central role in the adaptation to new diets, contributing to the efficient assimilation and detoxification of their food (Husnik and McCutcheon 2018; Prasad et al. 2021; Xia et al. 2021). Another interesting example of the importance of horizontally acquired genes in insects is the case of the *gasmin* gene, which is required for the lepidopteran *Spodoptera littoralis* to combat its natural enemies and infection (Di Lelio et al. 2019). Recently, we have reported the horizontal acquisition of Ribosome Inactivating Protein (RIPs, EC 3.2.2.22) encoding genes by some species of insects (Lapadula et al. 2017; Lapadula et al. 2020b).

RIPs are RNA *N*-glycosidases that irreversibly modify ribosomes through the depurination of an adenine residue in the conserved alpha-sarcin/ricin loop (SRL) of rRNA (Endo and Tsurugi 1988). This modification in a key component of the ribosomal elongation-cycle machinery prevents the binding of the elongation factor 2 to the ribosome, arresting protein synthesis (Nilsson and Nygard 1986). It is known that RIP encoding genes are found in plant, bacterial and fungal lineages (Lapadula and Ayub 2017). The recent exponential increase in database information has boosted the power of homology searches allowing for the discovery of new members of RIP genes family. Using this approach, for the first time we reported the presence of RIP genes in the metazoa kingdom (Lapadula et al. 2017; Lapadula et al. 2020b; Lapadula et al. 2013). Up to date, the taxonomic distribution of these genes in animals is very narrow, restricted to mosquitoes from the Culicinae subfamily (Lapadula et al. 2017) (including *Aedes aegypti*) and whiteflies from the Aleyrodidae family (including *Bemisia tabaci*) (Lapadula et al. 2020b). A combination of phylogeny and synteny analyses revealed that both groups of genes are derived from two independent HGT events, probably from bacterial and plant donors, respectively. Moreover, in both groups of insects, we showed that the RIP open reading frames show signatures of evolution under purifying (negative) selection, strongly suggesting that they positively impact the fitness of host organisms (Lapadula et al. 2017; Lapadula et al. 2020b). Recently, we have also demonstrated that two of the three RIP genes present in the *A. aegypti* genome are transcribed, and that their transcripts are polyadenylated (Lapadula et al. 2020a). Most importantly, the expression levels of these RIP genes are modulated across the developmental stages of mosquitoes (Lapadula et al. 2020a). By using transcriptomic data, the expression of these foreign genes could also be confirmed in *B. tabaci* (Lapadula et al. 2020b). Altogether, these data support the hypothesis that RIP genes have a physiological role in insects.

Although several members of the RIP protein family have been extensively studied at the biochemical level, their biological roles remain open to speculation. In some cases, activities against viruses, microbes or parasites have been postulated for plant RIPs (Peumans et al. 2001; Stirpe 2013; Zhu et al. 2018). Recently, RIPs from the symbiotic *Spiroplasma spp*. (class Mollicutes) have been suggested to play a defensive role in Drosophila species by preventing the development of parasitic nematodes and wasps (Ballinger and Perlman 2017; Ballinger and Perlman 2019; Hamilton et al. 2016). Based on these reports, we consider that RIP genes fulfill an important functional niche in insects that would be filled either from horizontally transferred genetic material or by a symbiotic interaction. The toxic nature of this protein family makes it possible to postulate the hypothesis that RIP genes play a defensive role in insects. In the present work we report the occurrence of a third independent HGT event supporting the recurrent acquisition of RIP genes by this mechanism in insects. After analyzing public transcriptomic data, we found a convergence in temporal and spatial expression of these independently acquired genes, which is compatible with expression patterns of immune effectors. Moreover, we identified depurinated sequences belonging to pathogens ribosomes that induce RIP expression in *A. aegypti*, strongly suggesting the mechanism through which these ribotoxins could carry out the protective role. In summary, we provide new evidence supporting the hypothesis that these foreign genes are new immune effector molecules of insects.

## Results

### Recurrent acquisition of RIP genes by Horizontal gene transfer in insects

Recent database searches focused on insects have led us to find new RIPs in the swede midge *Contarinia nasturtii* (named RIPCn1 and RIPCn2) and in the fungus gnat *Bradysia odoriphaga* (named RIPBo1 and RIPBo2). These flies of Diptera order belong to the Cecidomyiidae and Sciaridae sister families, respectively. All sequences were recognized by Pfam as RIPs. Similar to whiteflies, fly RIPs have a putative signal peptide in the N-terminal end and at least RIPCn2 harbor introns. Moreover, a sequence alignment of predicted proteins revealed that the five residues responsible to form the active site of RIPs were fully conserved for three of these toxins (**Supplementary fig. 1**). RIPBo2 showed a premature stop codon encoding for a truncated protein lacking three of these key residues. BLAST analyses of the genomics scaffolds (JAFDOW010000841, VYII01002082 and VYII01000852) harboring these genes showed that most of the encoding protein sequences surrounding the RIP genes yielded maximum scores with arthropod annotated proteins (**Supplementary Datafile 1**). In contrast, RIPs showed maximum amino acid sequence identity to bacterial homologs (around 36%), and lower sequence identity to the previously described RIPs from mosquitoes and whiteflies (around 24%).

The phylogenetic tree (**Fig. 1A, Supplementary fig. 2**) shows that new RIP encoding genes from Sciaridae and Cecidomyiidae are monophyletic (Transfer Boostrap Expectation; TBE = 0.66) and are embedded in a clade of bacterial sequences (TBE = 0.74). Moreover, metazoan (insect) RIP does not form a clade. This result supports a common origin for fly RIPs belonging to these sister families, but independent from whitefly and Diptera homologues. Furthermore, *C. nasturtii* and *B. odoriphaga* genes form sister clades, revealing that the gene duplication events took place after divergence of these families, yielding two different paralogues in each lineage. Moreover, the absence of RIP genes in other species with fully sequenced genome of Sciaroidea superfamily (*e*.*g. Bradysia coprophila, Sitodiplosis mosellana, Mayetiola destructor* and *Catotricha subobsoleta*) suggests the occurrence of gene loss events (**Supplementary fig. 3**), a commonly observed pattern in this protein family (Lapadula et al. 2013).

**Fig. 1.**
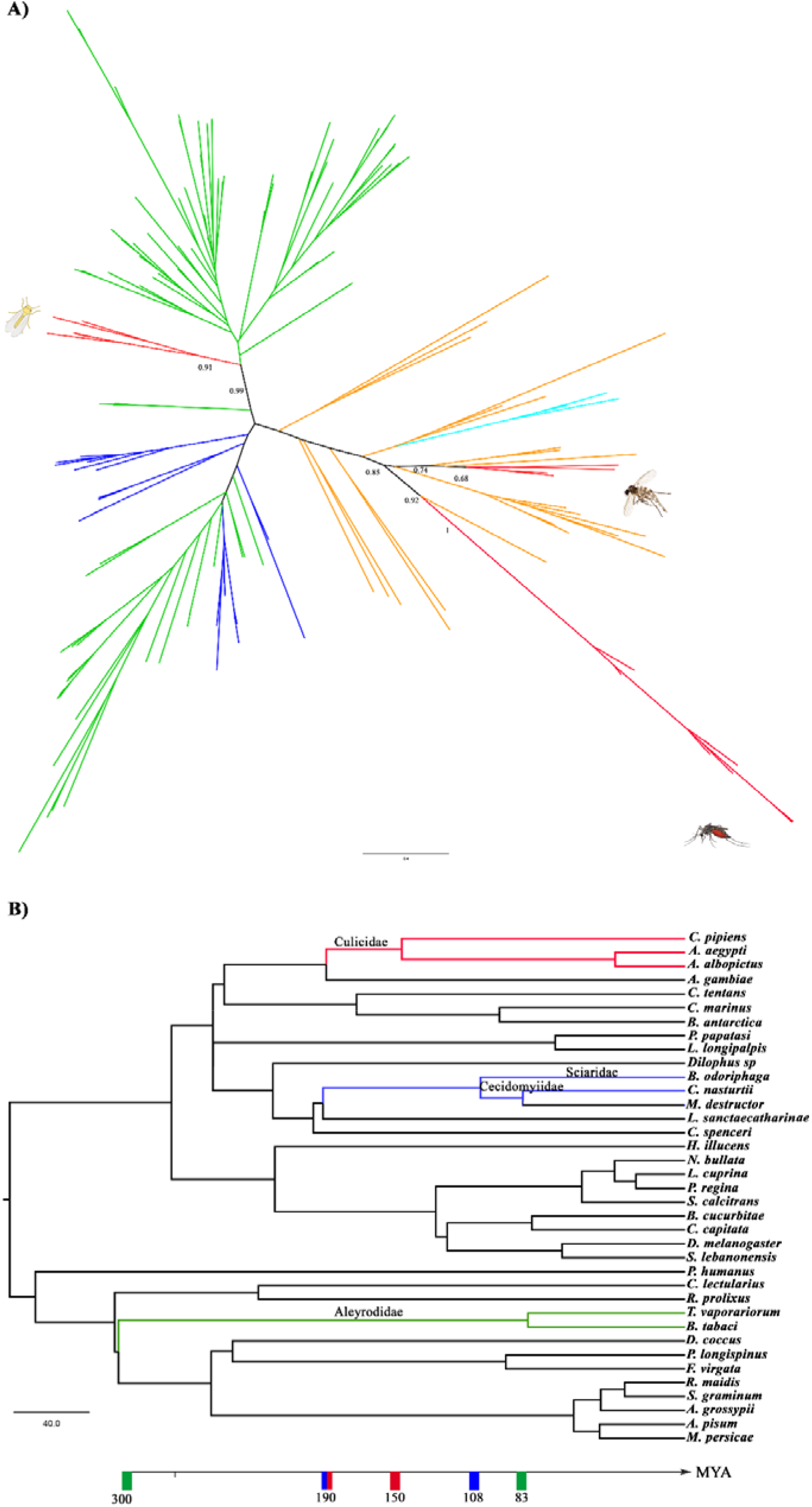
**A)** Unrooted phylogeny of RIP genes. Branches are colored according to taxonomy: bacteria (orange), Cyanobacteria (Cyan), plants (green), fungi (blue), metazoan (red). TBE support values of relevant divergences are shown at nodes. Fly, mosquito and whitefly clades are marked with silhouettes. Fully annotated phylogeny is available as Fig. Supplementary 2. **B)** Phylogeny of selected species from Neoptera orders. The tree including species from Diptera (24), Hemiptera (12) and Psocodea (1) orders with fully sequenced genomes was constructed with the TimeTree knowledge-base (Kumar et al. 2017). Insects harboring RIP genes are shown in red, blue and green branches. The occurrence time of three independent HGT events are graphically represented with the estimated time windows. Time in million years ago (MYA) is indicated at the bottom.

Hitherto, RIP genes have been found in three clades of insects (**Fig. 1B**). Previously, we proposed that whiteflies and mosquitoes acquired RIP genes by two independent HGT events from plants and bacteria, respectively (Lapadula et al. 2017; Lapadula et al. 2020b). In the case of whiteflies this event took place before the divergence of *B. tabaci* and *T. vaporariorum* species in the range of 300 and 83 MYA (**Fig. 1B**). In mosquitoes these genes were acquired between the divergence of Anopheles and Culex/Aedes lineages and before the separation of Aedes and Culex genus between 190 and 150 MYA (**Fig. 1B**).

According to the phylogeny (**Fig. 1A**), fly RIPs share a common origin but their history is independent from previously reported RIPs in insects. The homology searches using fly RIPs as queries in complete genomes of insects other than Sciaridae and Cecidomyiidae families did not retrieve any new hits. These results indicate that fly RIPs are not derived from vertical inheritance through the insect lineage or any species previously reported to have RIP genes. Therefore, the most parsimonious hypothesis explaining the presence of RIP genes in these sister families is a third HGT event. This acquisition took place in a range of 190 and 108 MYA after Sciaridae and Cecidomyiidae cenancestor diverged from the other families belonging to the Sciaroidea superfamily (**Fig. 1B**). The fact that fly RIPs are embedded in a clade of bacterial homologues (**Fig. 1A**) indicates that the most likely donor is a prokaryotic organism.

In summary, we found evidence of a third HGT event for RIP genes in insects. The recurrent acquisition by this evolutionary mechanism supports the hypothesis that members of the RIP family have found a functional niche in these organisms. In the following sections we show evidence that supports a convergence in transcription profiles of different insect lineages that independently acquired these ribotoxins encoding genes.

### RIPs genes show higher expression levels in early stage of insect ontogeny

Recently, we reported that two of the three RIP encoding genes present in *A. aegypti* are transcribed and their expression is modulated across the developmental stage (Lapadula et al. 2020a). In this work we found that RIPae2 expression was higher for L4 and pupal stages while RIPAe3 showed the highest expression values at L3 and L4 stages. From the analysis of transcriptome information available in BioProject PRJNA419241 (Matthews et al. 2018), we observed a similar expression pattern of RIPAe2 achieving maximal values for early pupal stage and RIPAe3 in L4 stage (**Fig. 2A**). The abundance of transcript in sister species *A. albopictus*, (harboring seven RIP genes) indicated that different paralogous genes have different expression profiles throughout their development (**Fig. 2B**). Genes RIPAl1 and RIPAl2 are expressed in adult males while RIPAl3 and RIPAl6 transcripts are found between L1 and pupal stages. As it was the case for *A. aegypti* in this species the highest expression levels for RIPs genes as a whole are found in early stages of ontogeny.

**Fig. 2.**
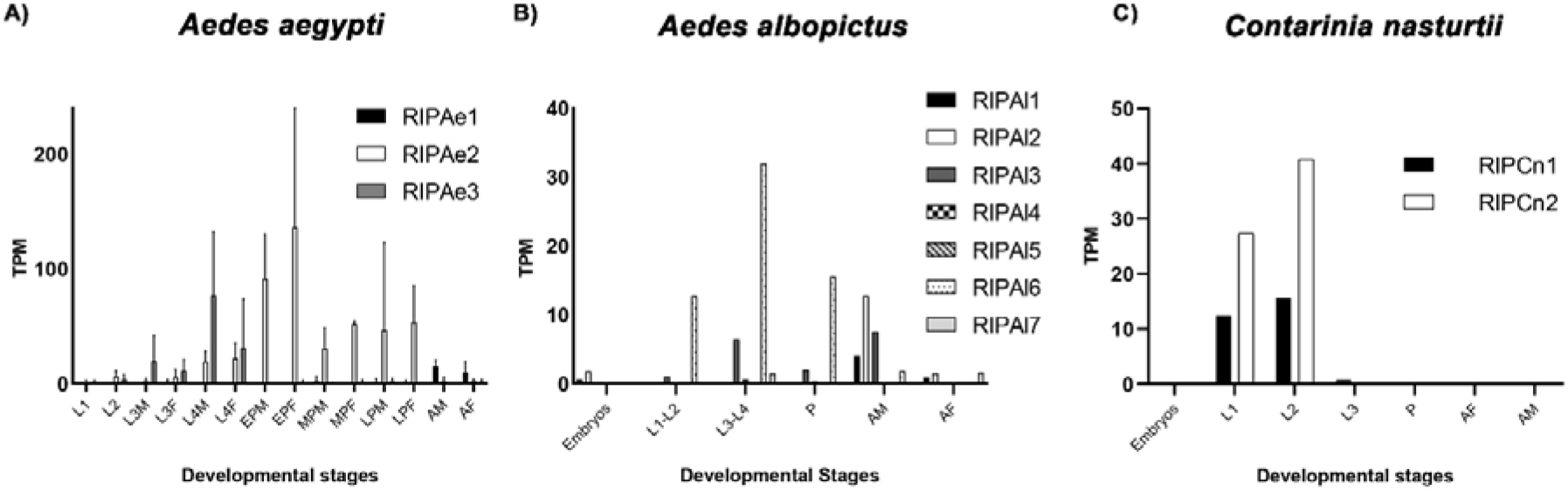
Expression of RIP genes across the developmental stages of insects as determined from transcriptomic assays. Expression of RIP genes was represented in transcript per millions (TPM). **A)** Larval 1-4 (L1-L4), early, mid and late pupal (EP, MP, LP) and adult stages (A) of *A. aegypti*. In some stages both sexes female (F) and male (M) were evaluated. Sequences Read Archives (SRA) files were taken from the BioProject PRJNA419241. **B)** Embryos, larval 1-2 (L1-2), larval 3-4 (L3-4), pupal (P), male adult (AM) and female adult (AF) of *A. albopictus*. SRA files were taken from BioProject PRJNA275727. **C)** Embryos, larval 1-3 (L1-L3), pupal and both sexes of adult stage (AF and AM) in *C. nasturtii*. SRA files were taken from the BioProject PRJNA646761.

In the case of *C. nasturtii* the abundance of transcripts obtained from BioProject PRJNA565761 showed that both RIPCn1 and RIPCn2 are expressed. According to this experiment we found the highest expression level for L1 and L2 while in other stages (embryos, L3, pupal and adults of both sexes) no transcripts of these genes were detected (**Fig. 2C**). In *B. odoriphaga* the absence of reference transcriptome prevented the building of the index to determine the abundance of transcripts. However, we performed an estimation by BLASTn searches. For this, we counted the number of retrieved hits after performing searches against Sequences Read Archives (SRA) files of BioProjects PRJNA388516 and PRJNA304774 using RIPBos as queries. This analysis indicated that the highest number of retrieved hits for RIPBo1 were between L2 and L4, followed by pupal stages (**Table 1**). Interestingly, RIPBo2 that encodes for a truncated variant showed no significant transcription in any stage. Thus, these results indicate that RIP encoding genes are modulated during insect ontogeny with a trend to the transcription during early development such as larval and pupal stages.

**Table 1.** Hits retrieved by BLASTn searches for SRA files of BioProjects PRJNA388516 and PRJNA304774. Mean of hits obtained for RIPBo1 and RIPBo2 in SRA files belonging to egg (E), larval 2 (L2), larval 4 (L4), pupal (P) and adult (A) stages are indicated in the second and third columns.

| Stage and SRA files | Mean of retrieved hits of PRJNA388516 (RIPBo1-RIPBo2) | Retrieved hits of PRJNA304774 (RIPBo1-RIPBo2) |
| --- | --- | --- |
| **E (SRX2879613/SRX2879612/SRX2879610)** | 1-0 | ND |
| **L2 (SRX2879611/SRX2879609/SRX2879608)** | 1700-3 | ND |
| **L3 (SRX1459738)** | ND | 1753-4 |
| **L4 (SRX2879607/SRX2879606/SRX2879604) (SRX1473238)** | 1500-4 | 2220-10 |
| **P (SRX2879616/SRX2879615/ SRX2879605) (SRX1473242)** | 365-1 | 151-0 |
| **A (SRX2879618 /SRX2879614/SRX2879617)** | 5-0 | ND |

ND\* non detected

### RIPs are expressed in body parts involved in immune response

In the previous section we described the expression profile of RIP genes along the ontogeny of insects. In order to determine those tissues and body parts where transcripts of these foreign genes are present, we performed a similar analysis in different transcriptomic experiments available in databases. If these genes are involved in immune response, it is expected that their transcript will be present in body parts like thorax or abdomen, where immune tissues such as fat body, gut or hemocytes are located. According to previously observed for adults (**Fig. 2A**), RIPAe2 is slightly expressed in the whole body of female *A. aegypti*. Despite this, from the analysis of Aegypti-Atlas, (Hixson et al. 2022) we found that their transcripts are mostly present in thorax and at a lower level in head and abdomen (**Fig. 3A**). Other body parts such as ovaries, Malpighian tubules and gut do not show the presence of these transcripts. The analysis of BioProject PRJNA236239 (Matthews et al. 2016) indicated that RIPAe1 is mostly expressed in maxillary palp of adult females, while in males, their expression is the highest in the abdominal tip (**Fig. 3B**). On the other hand, transcripts of RIPAe2 have similar expression levels for both sexes in body parts like rostrum, abdominal tip and brain (**Fig. 3B-C**). RIPAe3 is not detected in any body part for both sexes of adult insects. BioProjects PRJNA687261 (Romoli et al. 2021) and PRJNA548563 (Filosa et al. 2019) contain transcriptomic information of midgut and hindgut obtained from L3 and adult stages, respectively. In the L3 stage we observe that RIPAe2 and RIPAe3 transcripts are present in whole larva samples. However, in midgut these genes are not expressed indicating their absence in this tissue (**Supplementary fig. 4A**). On the other hand, RIPAe1 and RIPae2 were expressed in hindgut of adult individuals. Interestingly, their expression level is in the top quartile (**Supplementary fig. 4B-C**). Finally, in transcriptomes of Malpighi tubules obtained from BioProjects PRJNA246607 and PRJNA595990 no RIPs transcripts were detected.

**Fig. 3.**
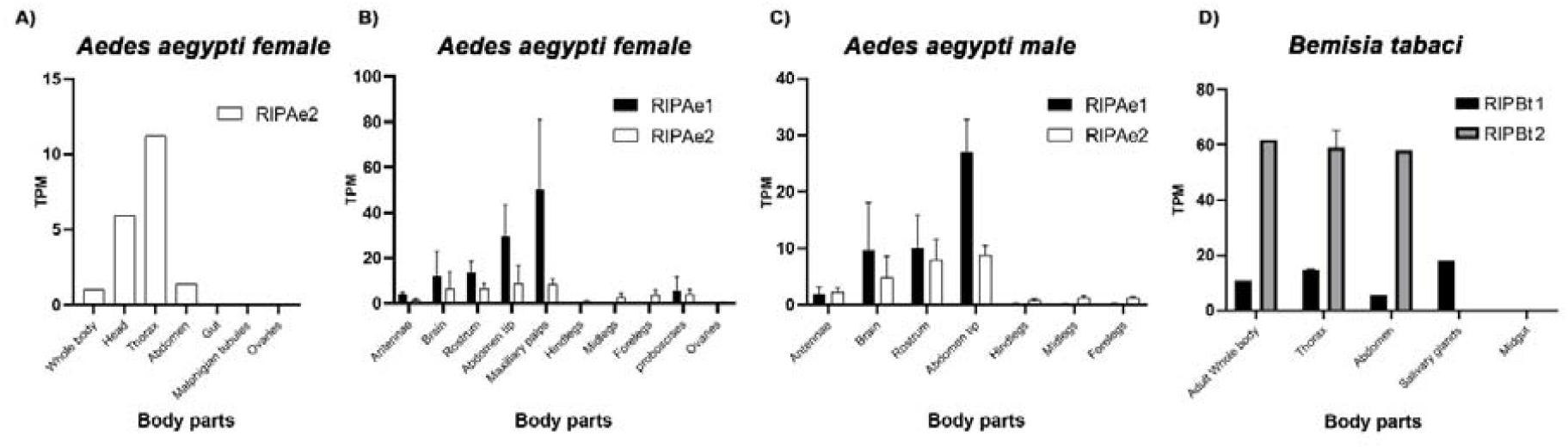
Expression of RIP genes in different body parts of insects. **A)** Expression of RIPAe2 gene from *A. aegypti* was represented in TPM for whole body, head, thorax, abdomen, gut, Malpighian tubules and ovaries of adult female. These results were taken from Aegypti-Atlas (Hixson et al. 2022) **B-C)** Expression of RIP genes were represented in TPM for adult females and males of *A. aegypti*, respectively. The expression was evaluated in antennae, brain, rostrum, abdominal tip, hindlegs, midleg and forelegs for both sexes while in female maxillary palp, proboscises and ovaries were included in the analysis. SRA files were taken from BioProject PRJNA236239. **D)** Expression of RIP genes of *B. tabaci* was represented in TPM for whole body, thorax, abdomen, salivary glands and midgut of adult individuals. SRA files were taken from the BioProject PRJEB26594.

Consistent with previously reported data of whiteflies *B. tabaci* (Lapadula et al. 2020b) transcripts of both RIP genes, RIPBt1 and RIPBt2, were found in the whole body of the adult stage, being the expression level of RIPBt2 higher than RIPBt1. Interestingly, transcripts of these genes are mostly found in thorax and abdomen while in salivary glands only RIPBt1 was expressed (**Fig. 3D**).

These results indicate that RIP transcripts are present in different body parts of insects. However, their presence is mostly located in abdomen and thorax of mosquitoes and whiteflies. Moreover, it was possible to identify expression signals in the hindgut of adult mosquitoes. Once again, we observed a convergence in expression profiles of these foreign genes in different lineages of insects. Although additional studies are needed to determine the exact location of RIP transcripts in abdomen and thorax, these analyses constitute a piece of evidence that support the presence of mRNA in body parts involved in the immune system.

### RIPs genes expression is increased after the infection with pathogens

If these foreign genes are immune effector molecules of insects, their transcription would be expected to be triggered after the infection with different pathogens. In order to find evidence supporting this hypothesis we analyzed information derived from bibliography for *A. aegypti*. In this species only RIPAe2 (AAEL008050) encoding gene has been annotated in VectorBase biasing bibliographic analysis. For this gene we found increased expression after the infection with several pathogens (bacteria, nematodes, and fungi) in adult mosquitoes. The most striking examples found was its upregulation post-infection with *Wolbachia pipientis* wMelPop (Kambris et al. 2009) and the Microsporidia *Edhazardia aedis* (Desjardins et al. 2015).

The analysis of RNAseq experiments after the infection with the nematode *Brugia malayi* (Choi et al. 2014; Juneja et al. 2015) in different refractory and susceptible strains of *A aegypti* support the potential of these genes as immune effector molecules in insects. Susceptible strains of mosquitoes support the development of nematode. On the contrary, in refractory strains parasites fail to develop and die within a few days. The transcriptome of whole body obtained from BioProject PRJNA255467 (Juneja et al. 2015) showed the upregulation of RIPAe2 for both strain (LVP-IB12^R^ and LVP-FR3^S^) of mosquitoes after the infection with the nematode (**Fig. 4A-B**). Moreover, the expression level of RIPAe2 increases along time, its highest being at 48 hours post infection. In this experiment, RIPAe1 showed no difference in its expression after the infection with the pathogen and their TPM values were always lower than RIPAe2. The RNA-seq experiment in thorax of *A. aegypti* obtained from BioProject PRJNA232599 (Choi et al. 2014) was consistent with the ones observed in the whole body. In blackeyed Liverpool (BEY-LVP) susceptible strain RIPae2 was significantly upregulated only at day 1 and 8 post infection with means of TPM values of 258 and 207, respectively (**Fig. 4F**). On the other hand, RIPae1 had lower expression level in this strain and it never showed differences between infected and uninfected conditions for the evaluated days (**Fig. 4E**). In the refractory *A. aegypti* (RED) strain both genes RIPAe1 and RIPae2 always showed higher levels of transcription after the treatment with the nematode (**Fig. 4C-D**). Moreover, the TPM values obtained for both genes in this strain were higher than the values observed in BEY-LVP strain, suggesting that RIPs might be involved in the resistance against the parasite. The highest TPM values observed in thorax for both genes support the hypothesis that their expression is enriched in this mosquito’s body part, which is consistent with data presented in **Fig. 3A**. In these experiments, no reads were detected for RIPae3 in any conditions, suggesting that its expression is not induced by pathogen infection in the adult stage. However, the transcriptome of L3 after the infection with *E. coli* (Romoli et al. 2021) indicated that RIPAe3 has an increase in level of transcription 20 hours after the infection (**Supplementary fig. 4A**). From the analysis of transcriptome experiments where virus infection is evaluated, no modulation in RIP expression was observed. Therefore, here we show evidence that RIPs expression is upregulated in *A. aegypti* after the infection with different pathogens like fungus, bacteria or nematodes. Moreover, the results presented suggest that RIP genes may contribute in defense against the nematode *B. malayi* in refractory strains of mosquitoes.

**Fig. 4.**
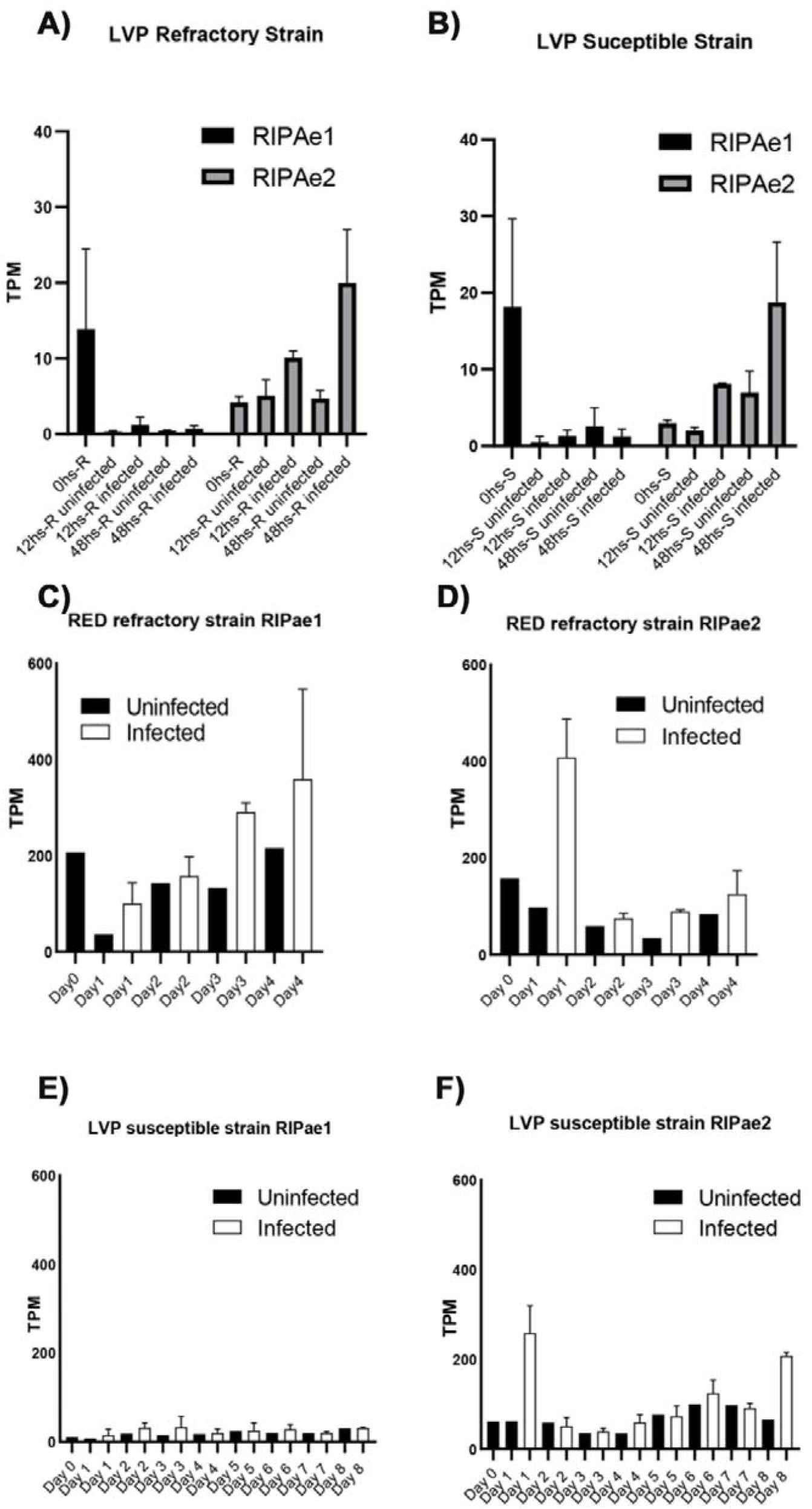
Expression of RIP genes in adults *A. aegypti* after the infection with the nematode *B. malayi*. **A-B)** Reads were taken from BioProject PRJNA255467 using the whole body of LVP-IB12^R^ and LVP-FR3^S^ strains, respectively. Expression of RIPAe1 and RIPAe2 genes was represented in TPM for different times after the infection. **C-F)** Reads were taken from BioProject PRJNA232599 corresponding to thorax tissue of refractory BEY-LVP (**C-D**) and susceptible RED (**E-F**) strains, respectively. Expression of RIPAe1 and RIPAe2 genes was represented in TPM for different times after the infection.

### Evidence of RNA *N*-glycosidase activity of *A. aegypti* RIPs

The toxicity of this family of proteins is presumably a consequence of their enzymatic activity. These toxins are RNA *N*-glycosidases that irreversibly modify ribosomes through the depurination of the first adenine residue in the G**<u>A</u>**GA motif present in the conserved SRL of rRNA. After the retrotranscription process, the reverse transcriptase preferentially inserts a dAMP opposite to the abasic site, which will result in a complementary dTMP after the first round of amplification step (**Fig. 5A**). Therefore, we searched for evidence of depurination in ribosomes of *B. malayi* from the PRJNA232599 transcriptomic experiment. For this we performed BLASTn searches against SRA files using a region of 61 bp from the 28S rRNA of nematode as query, including the SRL (**Fig. 5A**). Consistent with the report by (Choi et al. 2014), we observed that in susceptible BEY-LVP strain, the number of total reads increased during the course of infection, as expected from the nematode growth. Interestingly, we found few reads with a depurinated site between 4^th^ and 8^th^ days after the infection (**Fig. 5B**). In the refractory RED strain, the number of reads post infection in all samples is similar suggesting that the number of nematodes did not increase over time. In addition, in contrast with BEY-LVP, we detected reads with depurination signals in all the SRA files belonging to infected samples. Although in this strain the number of total retrieved reads was lower than for the BEY-LVP strain, the percentage of depurinated reads was higher, achieving 43% of retrieved sequences at day 4 (**Fig. 5B**). Thus, our results are the first piece of evidence supporting that these horizontally acquired genes of insects encode functional enzymes. On the other hand, these results support RNA *N*-glycosidases activity to be involved in defense response.

**Fig. 5.**
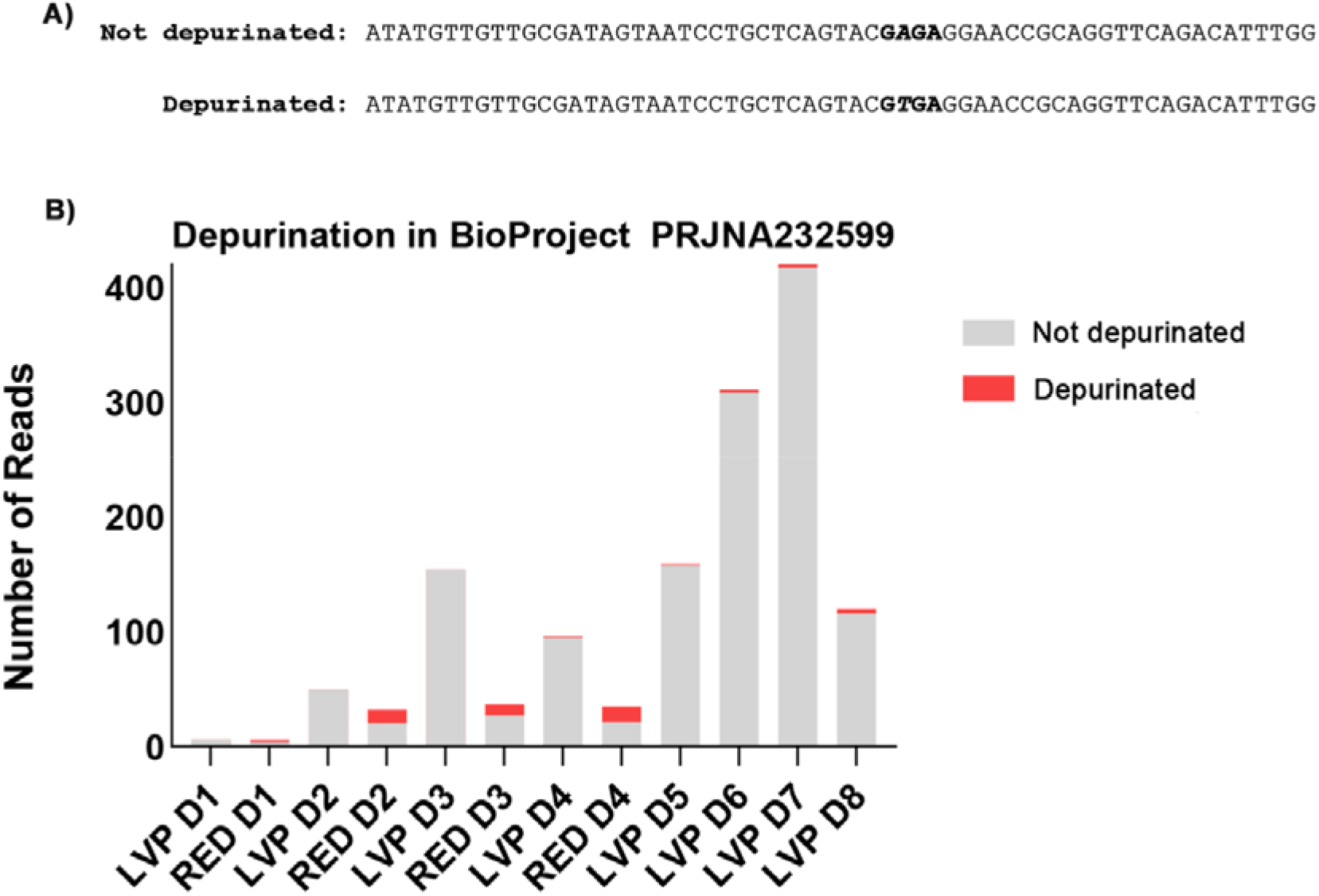
Evidence of RNA *N*-glycosidases activity. **A)** Not depurinated and depurinated sequences of 28S rRNA of *B. malayi* used to perform BLAST searches. The GAGA motif of the SRL is indicated with bold letters. The position which is target of depurination, adenine and thymine, are indicated with italic bold letters. **B)** Number of reads retrieved by BLAST searches in SRA files of infected samples from BioProject PRJNA232599. Not depurinated and depurinated reads are indicated with grey and red colors, respectively.

## Discussion

RIPs have been largely described in plant and bacterial lineages (Bolognesi et al. 2016; Di Maro et al. 2014; Stirpe 2004). In recent years, we have demonstrated the presence of RIP genes in fungi and metazoa (Lapadula and Juri Ayub 2017; Lapadula et al. 2020b; Lapadula et al. 2013). In animals, the presence of these ribotoxins is restricted to a few species of insects, being this narrow and patchy taxonomic distribution a hallmark of metazoan RIPs. The first RIP that we reported in insects belong to mosquitoes from the Culicinae subfamily (Lapadula et al. 2017; Lapadula et al. 2013). Then, we confirmed the presence of these genes in a second group belonging to whiteflies from the Aleyrodidae family (Lapadula et al. 2020b). Here, we found evidence of RIP encoding genes in a third lineage of insects belonging to the Sciaroidea superfamily. In this case these genes are present in two species of flies (*C. nasturtii* and *B. odoriphaga*). From the primary structure of genes, it was evident that all RIP encoding genes found in insects showed low sequence identity among them. Furthermore, features such as the paralogues number, the presence of signal peptides or introns (**Table 2**) differ for each lineage. Phylogenetic analysis (**Fig. 1**) indicated that metazoan RIPs does not form a monophyletic group. All these pieces of evidence support the idea that metazoan RIP genes have independent origins. Previously, we have reported that ribotoxins encoding genes were horizontally acquired by mosquitoes (Lapadula et al. 2017) and whiteflies (Lapadula et al. 2020b) from bacterial and plant donors, respectively. Here, we propose that flies RIPs were acquired by a third HGT event from a bacterial organism. The sister clade is shaped by sequences from the entomopathogen genera *Photorhabdus* and phytopathogens genus *Xanthomona* and *Brenneria* (**Supplementary fig. 2**). Across their ontogeny, organisms which belong to the Sciaroidea superfamily live in soil or host like fungus and plants. This superfamily includes fungivorous organism such as mycetophilids (Jakovlev 2012), and others like the Sciaridae family which live as larvae primarily in soil litter feeding on plant roots (Binns 1981) and Cecidomyiidae whose larvae produce secretions that dissolve the waxy cuticle and liquefy the underlying cells of the surrounding leaf surface (Readshaw 1966). This last indicated that the cenancestor of these flies were likely exposed to a large number of bacteria present in their habitats. It has been reported that HGT could be facilitated in early developmental stages of insects by the weakness of the Weisman barrier in these moments of their lifecycle (Huang 2013). The most likely donor are organisms sharing the same ecological niche such as symbiont (Dunning Hotopp et al. 2007; Kondo et al. 2002) or like -we previously proposed-plant and microbe-feeding insects (Lapadula et al. 2017; Lapadula et al. 2020b). This intimate association between recipient and donor organisms may facilitate the HGT.

**Table 2.**
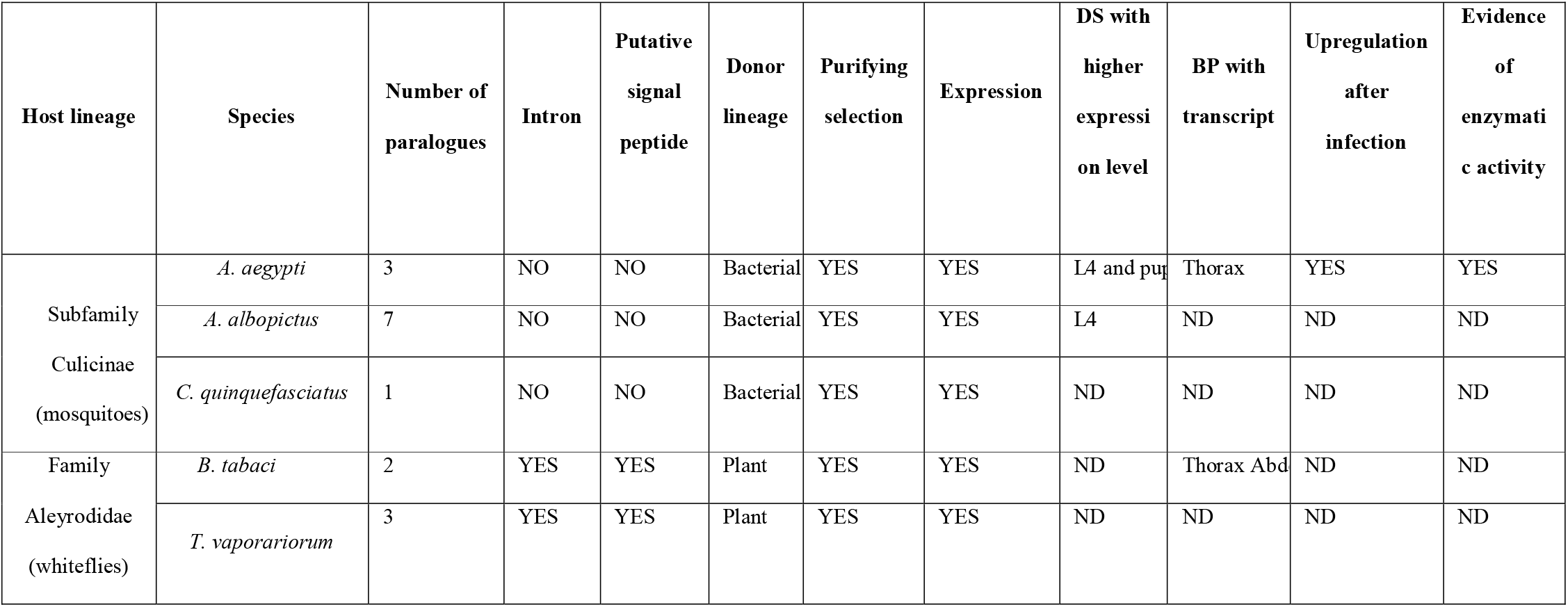
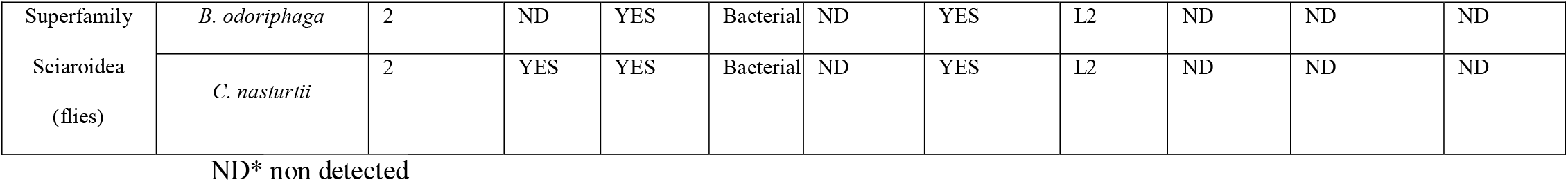
Information of RIP encoding genes in insects. Number of paralogues, presence of introns and signal peptides, potential donor lineage, evidence of evolution under purifying selection, evidence of expression, transcriptional information such as expression in developmental stages (DS), body parts (BP), post infection with pathogens and evidence of enzymatic activity are indicated for each specie.

Although horizontally transferred genes have been related to diet adaptation in several organism like arthropods (Wybouw et al. 2016) and insect (Prasad et al. 2021) there is also evidence that foreign genes could be involved in host defense (Di Lelio et al. 2019; Husnik and McCutcheon 2018). In insects some examples of HGT acquired genes involved in defense include bacterial lysozymes acquired by pea aphid, *Acyrthosiphon pisum* (Nikoh et al. 2010) and by *Halyomorpha halys* (Ioannidis et al. 2014). Recently, the acquisition by HGT of five toxin genes in *C. nasturtii* species was reported, which plays a nontrivial new role in insect immune function against eukaryotic enemies (Verster et al. 2021). In line with our model, the authors postulated that most likely donors are microbes that share the same environment with swede midge. The recurrent acquisition of RIP encoding genes by insects, the maintenance of these genes in host genomes by effects of natural selection and the occurrence of transcription are pieces of evidence that strongly support a functional role for these foreign genes in these organisms. The toxic nature of this protein family and the defensive roles that RIPs from *Spiroplasma* play in *Drosophila* species that lack these genes (Ballinger and Perlman 2017; Ballinger and Perlman 2019) further suggests that these ribotoxins could be immune molecules effectors.

Insect innate immunity includes both cellular and humoral responses (Ali Mohammadie Kojour et al. 2020; Hoffmann et al. 1996). The first is performed by hemocytes and consists predominantly of phagocytosis, nodulation and encapsulation of invading microorganisms (Ali Mohammadie Kojour et al. 2020; Hoffmann et al. 1996). The humoral response, mediated by soluble plasma proteins or fat body, implies clotting, melanin synthesis, and a rapid synthesis of a battery of antimicrobial peptides (AMPs) (Ali Mohammadie Kojour et al. 2020; Hoffmann et al. 1996). These molecules exhibit a broad spectrum of activity directed against bacteria and/or fungi. RIPAe2 is upregulated in response to infectious challenges caused by bacteria and fungi in *A. aegypti*, we also observed that the expression was triggered after the treatment with nematode *B. malayi*, we even observed depurination in nematode ribosome (**Fig.s 4 and 5**). Interestingly, (Choi et al. 2014) determined the host response profile comparing infected *vs*. uninfected *A. aegypti* BEY-LVP and found that genes with rRNA *N*-glycosylase activity (GO:0030598) are over-represented among the group of transcripts after the infection with *B. malayi*. In LVP-IB12^R^ and LVP-FR3^S^ strains (Juneja et al. 2015) reported a striking concordance between the transcriptional response of immune genes. Interestingly, they showed that RIPAe2 is one of the group of genes whose expression is induced in both strains after the infection (Juneja et al. 2015). *In vitro* evidence indicates that cecropin, an AMP from insect haemolymph, attenuates the motility of microfilariae of *Brugia pahangi* (Chalk et al. 1995). Thus, we postulate that RIP encoding genes could be immune effector molecules with regulation patterns comparable to AMPs.

The synthesis of AMPs is a hallmark of the host defense of higher insect orders like the Holometabola (essentially the Lepidoptera, Diptera, Hymenoptera and Coleoptera) and some Heterometabola (*e*.*g*. Hemiptera) (Hoffmann et al. 1996), organisms in which the presence of RIP encoding genes has been reported (**Table 2**). On the other hand, the presence of antibacterial activity and the expression of AMPs in early stages of insect ontogeny has been reported in some species. In the case of *Drosophila melanogaster*, a decrease in the number of bacteria at late pupal stage was reported, indicating antibacterial activity in this stage (Bakula 1969). Larvae of Black Soldier Fly, *H. illucens*, produced several AMPs which protect the insect from pathogens such as *E. coli* and *S. enterica* (Erickson et al. 2004). In a similar way the larvae of *Anopheles gambiae* mosquito, which live in a microbe-rich aquatic environment, exhibit higher levels of immune gene expression than adults (League et al. 2017). In *Bombix mori* at spinning and prepupa stages, a large increase in the expression of some AMPs was detected in the gut (Wu et al. 2010). The upregulation in the expression of AMPs in early stages of insect ontogeny is a consequence of hormonal regulations. 20-hydroxyecdysone is a steroid hormone produced by prothoracic glands prior to each moult. This hormone promotes humoral immunity by increasing the expression of AMP genes after immune challenge either by direct regulation or through interaction with other players of the immune response (Nunes et al. 2021). Transcriptomic experiments carried out in *A. aegypti, A. albopictus, B. odoriphaga* and *C. nasturtii* showed that the highest expression levels for RIPs genes are found in early stages of development (**Fig. 2 and Table 2**). These analyses were consistent with results previously reported in two strains of *A. aegypti* (Lapadula et al. 2020a). All this evidence supports a convergent expression profile across the ontogeny of insects.

In general, AMPs are produced after an immune challenge by the fat body that releases them into the haemolymph. These kinds of peptides are regulated at the transcriptional level, through the binding of the nuclear factor kappa-light-chain-enhancer of activated B cell (NF-κβ) (Manniello et al. 2021). RIPAe2 is found amongst the group of genes that are upregulated post-infection with *Plasmodium gallinaceum* in the fat body of transgenic *Ae. aegypti*, where the transcription factor NF-kB REL-2 is overexpressed (Zou et al. 2011). REL-1 and REL-2 are transcription factors that regulate the activation of genes downstream of the Toll and IMD pathways, the two main signaling cascades regulating insect immunity (Manniello et al. 2021; Valanne et al. 2011). In line with that, results shown here indicate that RIP transcripts are present in the thorax and abdomen of mosquitoes and whiteflies, body parts where the fat body is located (**Fig. 3 and Table 2**). Even in *A. aegypti* the level of transcripts present in thorax after the infection with *B. malayi* were higher than that found in the whole body (**Fig. 4**). These pieces of evidence indicate that RIP transcripts are present in body parts involved in immune response.

In plants, the RIP family has been associated in defense against several kinds of pathogens like fungus, bacteria, virus and insects (Zhu et al. 2018). In a similar way, the overexpression of RIPs reported in *A. aegypti* after the infection with the Microsporidia *Edhazardia aedis* (Desjardins et al. 2015), which allowed the authors to propose that rRNA *N*-glycosylase activity (GO:0030598) might play a role in the immune response of *A. aegypti*. On the other hand, bacterial RIPs from Spiroplasma endosymbiont are key in Drosophila defense against wasp (Ballinger and Perlman 2017; Ballinger and Perlman 2019; Hamilton et al. 2016). Here, we identified a higher number of depurinated sites on the SRL region that belong to a pathogen inducing RIP expression in *A. aegypti* refractory strain. Additional searches using the homologous region of 28S rRNA from mosquitoes as query did not retrieve any depurinated reads, suggesting that RIP genes do not have an effect on host ribosomes. The mechanisms proposed for resistance in mosquitoes against *B. malayi* include reduced ingestion of parasites, physical killing of parasites in the foregut, barriers to penetration of the midgut, and hemolymph factors that kill the parasite in the thoracic cavity and lead to melanotic encapsulation (Kobayashi et al. 1986). Therefore, the confirmation of the presence of depurination in ribosomes of *B. malagy* (**Fig. 5**) supports the hypothesis that these foreign genes have an impact on pathogen viability and contribute to immune response of infected organisms. RNA *N*-glycosidases activity could be the main mechanism through which these proteins play a defensive role in insects.

### Conclusion

In conclusion, although additional studies are needed, similarity in spatial and temporal expression profiles found in organisms where RIP encoding genes have been independently acquired support a functional convergence. Data from this study, along with previous information, prompted us to propose that RIPs are immune effector molecules in insects. This hypothesis is supported by the follow points:

I. The highest expression levels for these genes are found in early developmental stages of insects.
II. Transcripts of these genes are present in body parts involved in humoral immune response.
III. Transcription of RIP genes in *A. aegyti* is upregulated after the infection with several pathogens.
IV. These foreign proteins conserve their toxicity as a consequence of their enzymatic activity.
V. In refractory *A. aegypti* strain, the number of depurinated ribosome from *B. malayi* is higher than in samples of infected susceptible strains.

## Experimental Procedures

### Homology searches and sequence analyses

BLASTp homology searches were performed under default parameters on insect databases (excluding *Aedes, Culex, Bemisia*, and *Trialeurodes* genus) using a previously reported set of RIP sequences (Lapadula et al. 2020b) as queries. Bacterial sequences retrieved automatically annotated sequences from *Contarinia nasturtii* and *Bradysia odoriphaga* genome database. Then, these new sequences were used as queries in tBLASTn searches and new not annotated homologues were found in both species of flies. Pfam analysis was performed to confirm the presence of RIP domain (PF00161). The presence of signal peptide was predicted using SignalP 5.0 (https://services.healthtech.dtu.dk/service.php?SignalP-5.0). The presence of introns was analyzed, when available, comparing predicted mRNA with genomic DNA using the splign tool (https://www.ncbi.nlm.nih.gov/sutils/splign/splign.cgi). A full list of insect RIP encoding genes is available in **Supplementary Table 1**.

### Multiple sequence alignment and phylogenetic inferences

*C. nasturtii* and *B. odoriphaga* RIP amino acids sequences were added to our previously reported dataset of RIP (Lapadula et al. 2017; Lapadula et al. 2020b; Lapadula et al. 2013). This dataset was used for constructing a Multiple Sequences Alignment (MSA), as previously described (Lapadula et al. 2017; Lapadula et al. 2020b). This MSA containing 168 sequences and 159 residues was used to perform phylogenetic analysis by Maximum Likelihood in RAxML (version 8.2.10, available at https://github.com/stamatak/standard-RAxML) (Stamatakis 2014). The WAG substitution matrix was selected using ProtTest 3.4 (Darriba et al. 2011). To estimate the robustness of the phylogenetic inference, 500 rapid bootstrap (BS) were selected. Transfer bootstrap expectation was calculated in BOOSTER (Lemoine et al. 2018). Phylogenetic relationships and divergence times among species were obtained from TimeTree knowledge-base (Kumar et al. 2017). FigTree (version 1.4.2, available at https://tree.bio.ed.ac.uk/software/figtree) was used to visualize and edit the trees.

### Transcriptomic data analysis

BioProjects of transcriptome experiments carried out in *Aedes aegypti, Aedes albopictus, Bemisia tabaci* and *Contarinia nasturtii* were selected from the National Center for Biotechnology Information (NCBI) database (https://www.ncbi.nlm.nih.gov/). Sequence Read Archive (SRA) in FASTQ format were downloaded for the datasets (codes of SRA files used in this work are indicated in **Supplementary Tables 2-4**). The quality of each SRA file was evaluated using FastQC software (http://www.bioinformatics.babraham.ac.uk/projects/fastqc/). The abundance of transcripts from RNA-seq data was quantified using the Kallisto program (Bray et al. 2016) which estimated counts in Transcripts Per Millions (TPM). Not annotated RIP sequences of different species were incorporated in each reference transcriptomes obtained in FASTA format from the NCBI database. These files were used to build the index for each species in Kallisto. The TPM values obtained for RIP genes were represented in bar plots using GraphPad Prism version 5.00 for Windows. In the case of mosquito species databases such as VectorBase (https://vectorbase.org/vectorbase/app) and *Aegypti-Atlas* (http://aegyptiatlas.buchonlab.com/) were also analyzed.

### Quantification of SRL depurination

The depurination of the SRL by RIP RNA *N*-glycosidase activity yields an abasic site. Upon conversion, in a retrotranscription process, the reverse transcriptase preferentially inserts a dAMP opposite to the abasic site. Following the first round of amplification step, this yields a complementary dTMP. This is the basis used by (Pierce et al. 2011) to determine this enzymatic activity by qPCR. SRA files belonging to BioProject PRJNA232599 carried out in *Aedes aegypti* after the infection with *B. malayi* were used to quantify the number of reads derived from depurinated SRL. For this analysis, a region of 61 bp from the 28S rRNA of the pathogen including the adenine (not depurinated) and thymidine (depurinated) residues were used as queries in BLASTn searches. The searching parameters were set in order to exclude *A. aegypti* as a result. All retrieved reads for each SRA file were downloaded and aligned with MAFFT online server. In each alignment, the number of adenine and thymine present in the target position of depurination was counted and represented in a bar plot using GraphPad Prism version 5.00 for Windows.

## Acknowledgements

W.J.L and M.J.A. are members of the scientific career of CONICET. This work has been funded by grants from CONICET (Consejo Nacional de Investigaciones Científicas y Técnicas) PIP 2732, PIBAA 28720210100081CO to M.J.A and W.J.L and UNSL (Universidad Nacional de San Luis) PROICO 02-1720 to M.J.A. We would also thank Carolina Mirallas from the GAECI (English Language Writing Advise) for their linguistic corrections of an early version of this work. Finally, W.J.L thanks Pablo V. Lapadula, who died as a consequence of SARS-CoV-2, for always encouraging him to continue this research.

## Notes

**Conflict of interest** disclosure The authors declare that they have no conflict of interest.

### Competing Interest Statement

The authors have declared no competing interest.

## Reference

Ali Mohammadie Kojour, M, Han, YS and Jo, YH (2020) An overview of insect innate immunity. Entomological research 50: 282–291.

Bakula, M (1969) The persistence of a microbial flora during postembryogenesis of Drosophila melanogaster. Journal of invertebrate pathology 14: 365–374.

Ballinger, MJ and Perlman, SJ (2017) Generality of toxins in defensive symbiosis: Ribosome-inactivating proteins and defense against parasitic wasps in Drosophila. PLoS Pathog 13: e1006431.

Ballinger, MJ and Perlman, SJ (2019) The defensive Spiroplasma. Curr Opin Insect Sci 32: 36–41.

Binns, E (1981) Fungus gnats (Diptera: Mycetophilidae/Sciaridae) and the role of mycophagy in soil: a review.

Bolognesi, A, Bortolotti, M, Maiello, S, Battelli, MG and Polito, L (2016) Ribosome-Inactivating Proteins from Plants: A Historical Overview. Molecules 21.

Bray, NL, Pimentel, H, Melsted, P and Pachter, L (2016) Near-optimal probabilistic RNA-seq quantification. Nat Biotechnol 34: 525–7.

Chalk, R, Townson, H and Ham, PJ (1995) Brugia pahangi: the effects of cecropins on microfilariae in vitro and in Aedes aegypti. Exp Parasitol 80: 401–6.

Choi, YJ, Aliota, MT, Mayhew, GF, Erickson, SM and Christensen, BM (2014) Dual RNA-seq of parasite and host reveals gene expression dynamics during filarial worm-mosquito interactions. PLoS Negl Trop Dis 8: e2905.

Darriba, D, Taboada, GL, Doallo, R and Posada, D (2011) ProtTest 3: fast selection of best-fit models of protein evolution. Bioinformatics 27: 1164–5.

Desjardins, CA, Sanscrainte, ND, Goldberg, JM, Heiman, D, Young, S, Zeng, Q, Madhani, HD, Becnel, JJ and Cuomo, CA (2015) Contrasting host-pathogen interactions and genome evolution in two generalist and specialist microsporidian pathogens of mosquitoes. Nat Commun 6: 7121.

Di Lelio, I, Illiano, A, Astarita, F, Gianfranceschi, L, Horner, D, Varricchio, P, Amoresano, A, Pucci, P, Pennacchio, F and Caccia, S (2019) Evolution of an insect immune barrier through horizontal gene transfer mediated by a parasitic wasp. PLoS Genet 15: e1007998.

Di Maro, A, Citores, L, Russo, R, Iglesias, R and Ferreras, JM (2014) Sequence comparison and phylogenetic analysis by the Maximum Likelihood method of ribosome-inactivating proteins from angiosperms. Plant Mol Biol 85: 575–88.

Dunning Hotopp, JC, Clark, ME, Oliveira, DC, Foster, JM, Fischer, P, Munoz Torres, MC, Giebel, JD, Kumar, N, Ishmael, N, Wang, S, Ingram, J, Nene, RV, Shepard, J, Tomkins, J, Richards, S, Spiro, DJ, Ghedin, E, Slatko, BE, Tettelin, H and Werren, JH (2007) Widespread lateral gene transfer from intracellular bacteria to multicellular eukaryotes. Science 317: 1753–6.

Endo, Y and Tsurugi, K (1988) The RNA N-glycosidase activity of ricin A-chain. The characteristics of the enzymatic activity of ricin A-chain with ribosomes and with rRNA. J Biol Chem 263: 8735–9.

Erickson, MC, Islam, M, Sheppard, C, Liao, J and Doyle, MP (2004) Reduction of Escherichia coli O157:H7 and Salmonella enterica serovar Enteritidis in chicken manure by larvae of the black soldier fly. J Food Prot 67: 685–90.

Filosa, JN, Berry, CT, Ruthel, G, Beverley, SM, Warren, WC, Tomlinson, C, Myler, PJ, Dudkin, EA, Povelones, ML and Povelones, M (2019) Dramatic changes in gene expression in different forms of Crithidia fasciculata reveal potential mechanisms for insect-specific adhesion in kinetoplastid parasites. PLoS Negl Trop Dis 13: e0007570.

Goldenfeld, N and Woese, C (2007) Biology’s next revolution. Nature 445: 369.

Hamilton, PT, Peng, F, Boulanger, MJ and Perlman, SJ (2016) A ribosome-inactivating protein in a Drosophila defensive symbiont. Proc Natl Acad Sci U S A 113: 350–5.

Hixson, B, Bing, XL, Yang, X, Bonfini, A, Nagy, P and Buchon, N (2022) A transcriptomic atlas of Aedes aegypti reveals detailed functional organization of major body parts and gut regional specializations in sugar-fed and blood-fed adult females. Elife 11.

Hoffmann, JA, Reichhart, JM and Hetru, C (1996) Innate immunity in higher insects. Curr Opin Immunol 8: 8–13.

Huang, J (2013) Horizontal gene transfer in eukaryotes: the weak-link model. Bioessays 35: 868–75.

Husnik, F and McCutcheon, JP (2018) Functional horizontal gene transfer from bacteria to eukaryotes. Nat Rev Microbiol 16: 67–79.

Ioannidis, P, Lu, Y, Kumar, N, Creasy, T, Daugherty, S, Chibucos, MC, Orvis, J, Shetty, A, Ott, S and Flowers, M (2014) Rapid transcriptome sequencing of an invasive pest, the brown marmorated stink bug Halyomorpha halys. BMC genomics 15: 1–22.

Jakovlev, J (2012) Fungal hosts of mycetophilids (Diptera: Sciaroidea excluding Sciaridae): a review. Mycology 3: 11–23.

Juneja, P, Ariani, CV, Ho, YS, Akorli, J, Palmer, WJ, Pain, A and Jiggins, FM (2015) Exome and transcriptome sequencing of Aedes aegypti identifies a locus that confers resistance to Brugia malayi and alters the immune response. PLoS Pathog 11: e1004765.

Kambris, Z, Cook, PE, Phuc, HK and Sinkins, SP (2009) Immune activation by life-shortening Wolbachia and reduced filarial competence in mosquitoes. Science 326: 134–6.

Keeling, PJ and Palmer, JD (2008) Horizontal gene transfer in eukaryotic evolution. Nat Rev Genet 9: 605–18.

Kobayashi, M, Ogura, N and Yamamoto, H (1986) Studies on filariasis VIII: Histological observation on the abortive development of Brugia malayi larvae in the thoracic muscles of the mosquitoes, Armigeres subalbatus. Medical Entomology and Zoology 37: 127–132.

Kondo, N, Nikoh, N, Ijichi, N, Shimada, M and Fukatsu, T (2002) Genome fragment of Wolbachia endosymbiont transferred to X chromosome of host insect. Proc Natl Acad Sci U S A 99: 14280–5.

Kumar, S, Stecher, G, Suleski, M and Hedges, SB (2017) TimeTree: A Resource for Timelines, Timetrees, and Divergence Times. Mol Biol Evol 34: 1812–1819.

Lapadula, WJ and Juri Ayub, MJ (2017) Ribosome Inactivating Proteins from an evolutionary perspective. Toxicon 136: 6–14.

Lapadula, WJ, Marcet, PL, Mascotti, ML, Sanchez-Puerta, MV and Juri Ayub, M (2017) Metazoan Ribosome Inactivating Protein encoding genes acquired by Horizontal Gene Transfer. Scientific Reports 7: 1863.

Lapadula, WJ, Marcet, PL, Taracena, ML, Lenhart, A and Juri Ayub, M (2020a) Characterization of horizontally acquired ribotoxin encoding genes and their transcripts in Aedes aegypti. Gene 754: 144857.

Lapadula, WJ, Mascotti, ML and Juri Ayub, M (2020b) Whitefly genomes contain ribotoxin coding genes acquired from plants. Scientific Reports 10: 15503.

Lapadula, WJ, Sanchez Puerta, M. and Juri Ayub, M (2013) Revising the taxonomic distribution, origin and evolution of ribosome inactivating protein genes. PLoS One 8: e72825.

League, GP, Estevez-Lao, TY, Yan, Y, Garcia-Lopez, VA and Hillyer, JF (2017) Anopheles gambiae larvae mount stronger immune responses against bacterial infection than adults: evidence of adaptive decoupling in mosquitoes. Parasit Vectors 10: 367.

Lemoine, F, Domelevo Entfellner, JB, Wilkinson, E, Correia, D, Davila Felipe, M. De Oliveira, T and Gascuel, O (2018) Renewing Felsenstein’s phylogenetic bootstrap in the era of big data. Nature 556: 452–456.

Lerminiaux, NA and Cameron, ADS (2019) Horizontal transfer of antibiotic resistance genes in clinical environments. Can J Microbiol 65: 34–44.

Manniello, MD, Moretta, A, Salvia, R, Scieuzo, C, Lucchetti, D, Vogel, H, Sgambato, A and Falabella, P (2021) Insect antimicrobial peptides: potential weapons to counteract the antibiotic resistance. Cell Mol Life Sci 78: 4259–4282.

Matthews, BJ, Dudchenko, O, Kingan, SB, Koren, S, Antoshechkin, I, Crawford, JE, Glassford, WJ, Herre, M, Redmond, SN, Rose, NH, Weedall, GD, Wu, Y, Batra, SS, Brito-Sierra, CA, Buckingham, SD, Campbell, CL, Chan, S, Cox, E, Evans, BR, Fansiri, T, Filipovic, I, Fontaine, A, Gloria-Soria, A, Hall, R, Joardar, VS, Jones, AK, Kay, RGG, Kodali, VK, Lee, J, Lycett, GJ, Mitchell, SN, Muehling, J, Murphy, MR, Omer, AD, Partridge, FA, Peluso, P, Aiden, AP, Ramasamy, V, Rasic, G, Roy, S, Saavedra-Rodriguez, K, Sharan, S, Sharma, A, Smith, ML, Turner, J, Weakley, AM, Zhao, Z, Akbari, OS, Black, WCt, Cao, H, Darby, AC, Hill, CA, Johnston, JS, Murphy, TD, Raikhel, AS, Sattelle, DB, Sharakhov, IV, White, BJ, Zhao, L, Aiden, EL, Mann, RS, Lambrechts, L, Powell, JR, Sharakhova, MV, Tu, Z, Robertson, HM, McBride, CS, Hastie, AR, Korlach, J, Neafsey, DE, Phillippy, AM and Vosshall, LB (2018) Improved reference genome of Aedes aegypti informs arbovirus vector control. Nature 563: 501–507.

Matthews, BJ, McBride, CS, DeGennaro, M, Despo, O and Vosshall, LB (2016) The neurotranscriptome of the Aedes aegypti mosquito. BMC Genomics 17: 32.

Nikoh, N, McCutcheon, JP, Kudo, T, Miyagishima, SY, Moran, NA and Nakabachi, A (2010) Bacterial genes in the aphid genome: absence of functional gene transfer from Buchnera to its host. PLoS Genet 6: e1000827.

Nilsson, L and Nygard, O (1986) The mechanism of the protein-synthesis elongation cycle in eukaryotes. Effect of ricin on the ribosomal interaction with elongation factors. Eur J Biochem 161: 111–7.

Nunes, C, Sucena, E and Koyama, T (2021) Endocrine regulation of immunity in insects. FEBS J 288: 3928–3947.

Peumans, WJ, Hao, Q and Van Damme, EJ (2001) Ribosome-inactivating proteins from plants: more than RNA N-glycosidases? FASEB J 15: 1493–506.

Pierce, M, Kahn, JN, Chiou, J and Tumer, NE (2011) Development of a quantitative RT-PCR assay to examine the kinetics of ribosome depurination by ribosome inactivating proteins using Saccharomyces cerevisiae as a model. Rna 17: 201–210.

Prasad, A, Chirom, O and Prasad, M (2021) Insect herbivores benefit from horizontal gene transfer. Trends Plant Sci 26: 1096–1097.

Readshaw, J (1966) The ecology of the swede midge, Contarinia nasturtii (Kieff.)(Diptera, Cecidomyiidae). I.—Life-history and influence of temperature and moisture on development. Bulletin of Entomological Research 56: 685–700.

Romoli, O, Schonbeck, JC, Hapfelmeier, S and Gendrin, M (2021) Production of germ-free mosquitoes via transient colonisation allows stage-specific investigation of host-microbiota interactions. Nat Commun 12: 942.

Stamatakis, A (2014) RAxML version 8: a tool for phylogenetic analysis and post-analysis of large phylogenies. Bioinformatics 30: 1312–3.

Stirpe, F (2004) Ribosome-inactivating proteins. Toxicon 44: 371–83.

Stirpe, F (2013) Ribosome-inactivating proteins: from toxins to useful proteins. Toxicon 67: 12–6.

Valanne, S, Wang, JH and Ramet, M (2011) The Drosophila Toll signaling pathway. J Immunol 186: 649–56.

Van Etten, J and Bhattacharya, D (2020) Horizontal Gene Transfer in Eukaryotes: Not if, but How Much? Trends Genet 36: 915–925.

Verster, KI, Tarnopol, RL, Akalu, SM and Whiteman, NK (2021) Horizontal Transfer of Microbial Toxin Genes to Gall Midge Genomes. Genome Biol Evol 13.

Wu, S, Zhang, X, He, Y, Shuai, J, Chen, X and Ling, E (2010) Expression of antimicrobial peptide genes in Bombyx mori gut modulated by oral bacterial infection and development. Dev Comp Immunol 34: 1191–8.

Wybouw, N, Pauchet, Y, Heckel, DG and Van Leeuwen, T (2016) Horizontal gene transfer contributes to the evolution of arthropod herbivory. Genome Biol Evol.

Xia, J, Guo, Z, Yang, Z, Han, H, Wang, S, Xu, H, Yang, X, Yang, F, Wu, Q, Xie, W, Zhou, X, Dermauw, W, Turlings, TCJ and Zhang, Y (2021) Whitefly hijacks a plant detoxification gene that neutralizes plant toxins. Cell 184: 1693–1705 e17.

Zhu, F, Zhou, YK, Ji, ZL and Chen, XR (2018) The Plant Ribosome-Inactivating Proteins Play Important Roles in Defense against Pathogens and Insect Pest Attacks. Front Plant Sci 9: 146.

Zou, Z, Souza-Neto, J, Xi, Z, Kokoza, V, Shin, SW, Dimopoulos, G and Raikhel, A (2011) Transcriptome analysis of Aedes aegypti transgenic mosquitoes with altered immunity. PLoS Pathog 7: e1002394.

