## Supporting information for "Foreign Ribosome Inactivating Proteins as immune effectors in insects"

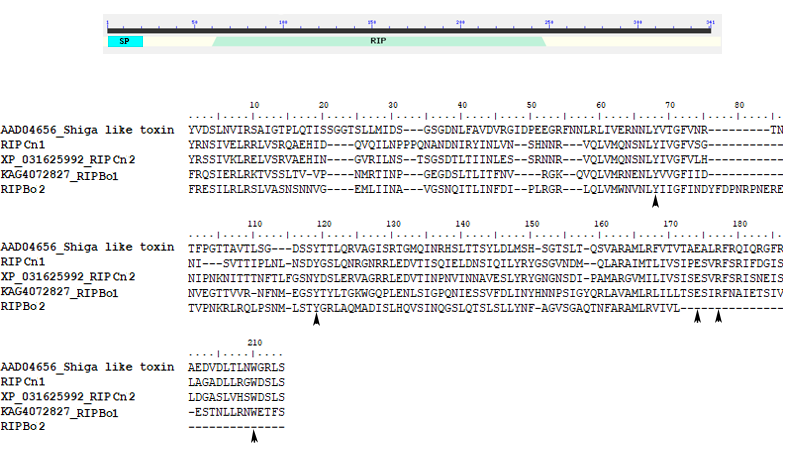

**Supplementary Fig. 1. Sequence analysis of RIPs from Sciaroidea superfamily. A)** Schematic representation of RIPBo1 protein sequence. The predicted protein harbors 341 amino acids. The predicted signal peptide (SP) and RIP domain regions are shown in light blue and green, respectively. **B)** Sequence alignment of the conserved RIP region from RIPCns and RIPBos along with SLT-1. Arrowheads indicate those residues predicted to form the active site.

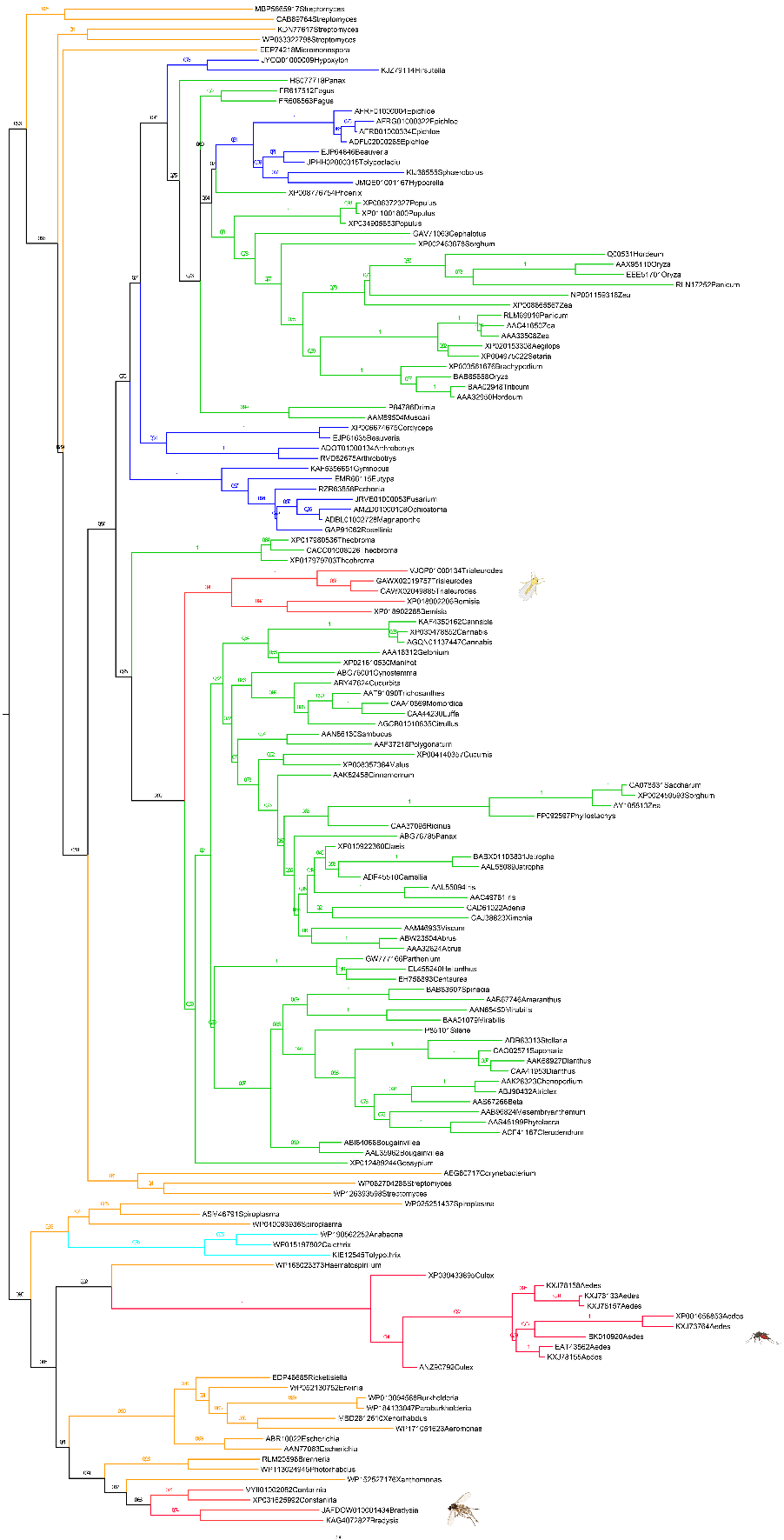

**Supplementary Fig. 2. RIPs fully annotated phylogeny.** Midpoint rooted phylogeny of RIP sequences. Tree was constructed in RAXML 8.2.12. Branches are colored according to taxonomy: bacteria (orange), cyanobacteria (cyan), plants (green), fungi (blue), metazoan (red). 500 rapid bootstrapping were performed and then transformed to TBE in BOOSTER. TBE support values are shown in nodes. Genus and GenBank accession codes are given for each taxa.

Until now, all the species with sequenced genomes into Aleyrodidae family and Culicinae subfamily have RIP genes. However, species with fully sequenced genomes as *Bradysia coprophila* belonging to Sciaridae and the species *Sitodiplosis mosellana*, *Mayetiola destructor* and *Catroticha subobsoleta* from Cecidomyiidae lack of RIP genes. In light of RIP phylogeny, we postulate that these genes have been vertically inherited from the cenancestor of both families, being later purged from other species genomes by gene loss events. In order to find evidence supporting this hypothesis we performed an analysis of synteny for Sciaridae and Cecidomyiidae species. Although in Bradysia species both genomics regions were syntenic the loss of RIP and other neighboring genes were confirmed for *B. coprophila*. This event is a consequence of duplications followed by a non-reciprocal translocation which removes the region where RIP is located. In the case of the Cecidomyiidae family only the homologous regions of RIPCn1 could be found in other species. RIP gene is the only sequence belonging to this contig absent in the other genomes.

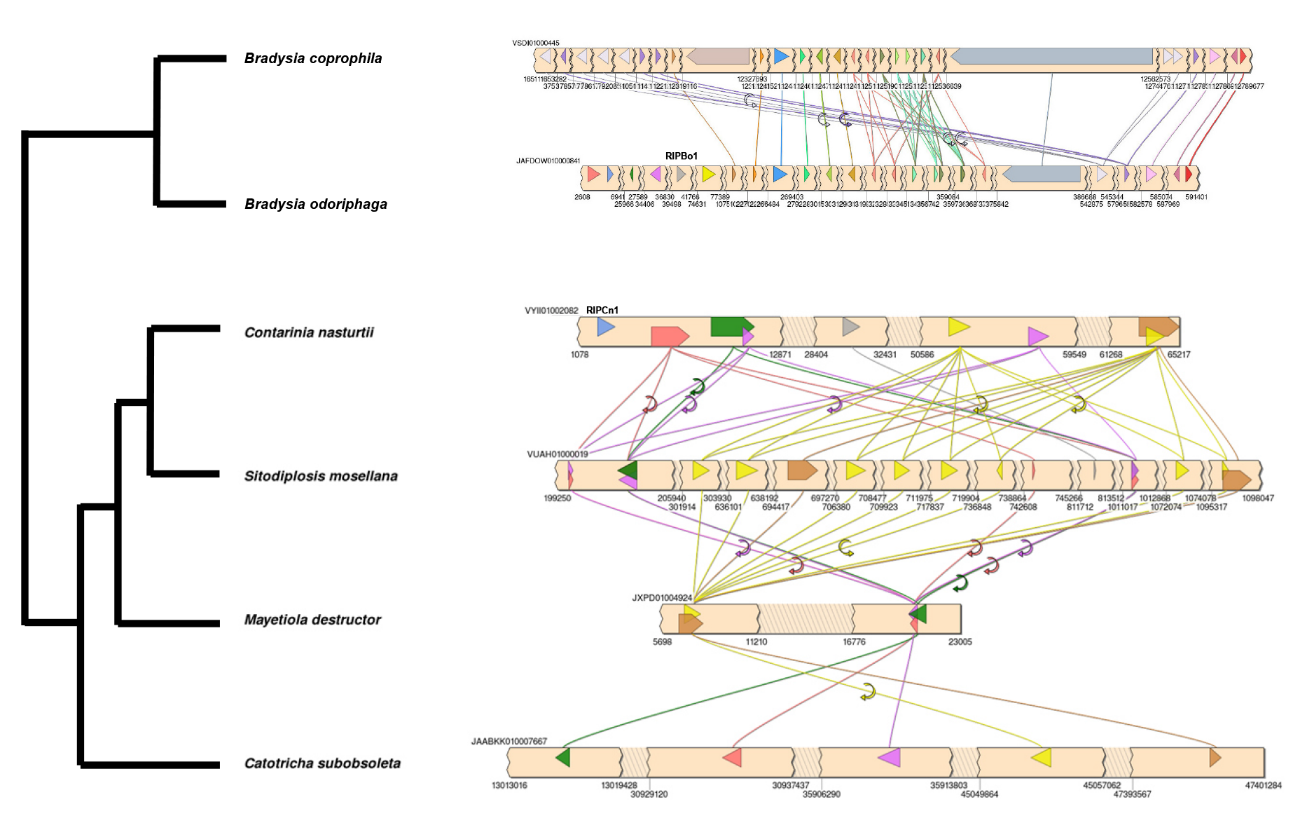

**Supplementary Fig. 3.** Scheme depicting the synteny of genomics regions harboring RIPBo1 and RIPCn1 genes. The contig containing RIP gene from *B. odoriphaga* (JAFDOW010000641) was compared with homologous region in *B. Coprophila* (VSDI01000445). The contig containing RIP gene from *C. nasturtii* (VYII01002082) was compared with homologous regions in *S. mosellana* (VUAH01000019), *M. destructor* (JXPD01004924) and *C. subobsoleta* (JAABKK010007667). Phylogenetic relationships among species were taken from a previous report^1^. Open reading frames are indicated with colored arrows. Regions corresponding to RIPBo1 and RIPCn1 are indicated on the respective arrows.

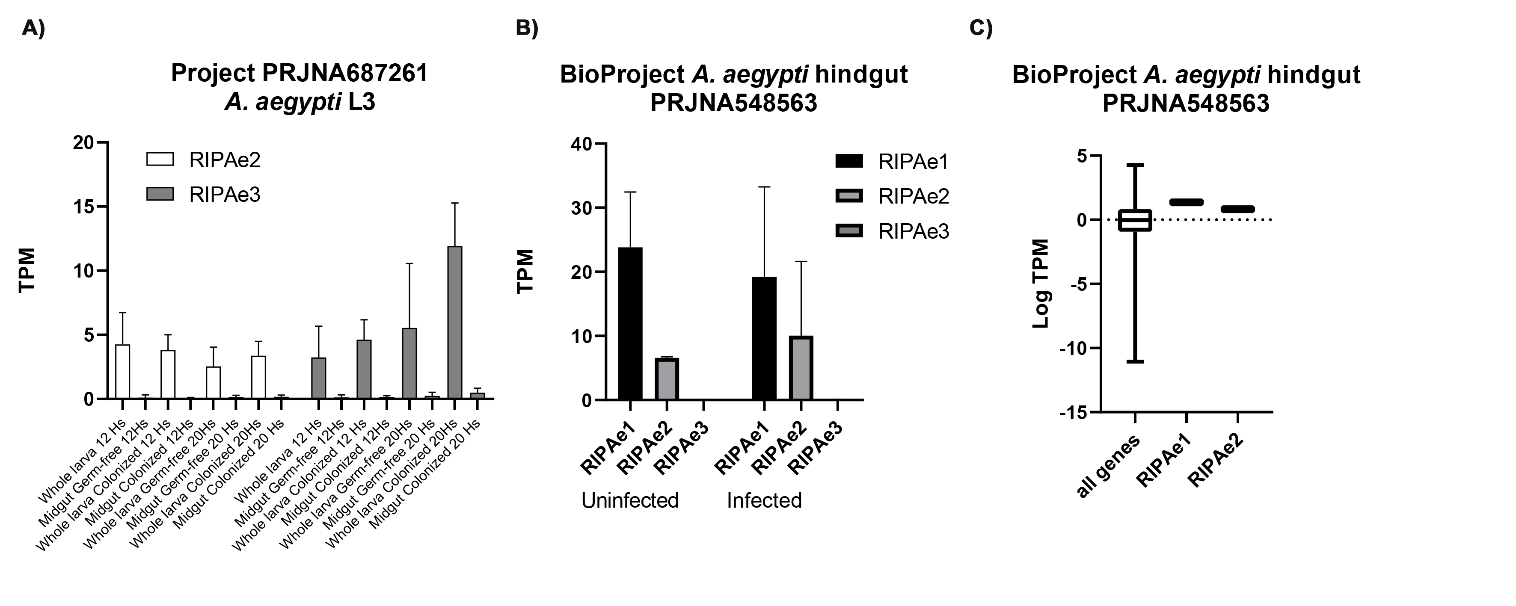

**Supplementary Fig. 4. Expression of RIP genes in midgut and hindgut of L3 and adult *A. aegypti* mosquitoes. A)** Expression of RIP genes of *A. aegypti* was represented in TPM for whole larva and midgut in organisms uninfected and infected with *Escherichia coli*. SRA files were taken from the BioProject PRJNA687261 **B)** Expression of RIP genes were represented in TPM for the hindgut of adults uninfected and infected with *Crithidia fasciculata*. SRA files were taken from the BioProject PRJNA548563. **C)** Mean of TPM values obtained for uninfected samples from the BioProject PRJNA548563 were plotted as log of TPM. Mean values of RIPAe1 and RIPAe2 were compared with the whole set of *A. aegypti* genes. The box-and-whisker plot shows expression levels as quartiles.

**Supplementary table 1.** Species of insects harboring RIP genes are indicated in the first column. RIP names used in this manuscript file are indicated in the second column. GenBank and VectorBase accession numbers of nucleotides and amino acids sequences are presented in the third and fourth columns, respectively.

| **Organism** | **RIP name** | **Nucleotide RIP codes (GenBank/VectorBase)** | **Amino acids RIP codes (GenBank)** |
| --- | --- | --- | --- |
| *Aedes aegypti* | RIPAe1 | MN299051 | EAT43562 |
|  | RIPAe2 | XM_001658803/**AAEEL008050** | XP_001658853 |
|  | RIPAe3 | BK010920 | DAC84270 |
| *Aedes albopictus* | RIPAl1 | **AALF005486** | KXJ78155 |
|  | RIPAl2 | **AALF005485** | KXJ78156 |
|  | RIPAl3 | **AALF005487** | KXJ78157 |
|  | RIPAl4 | **AALF005488** | KXJ78158 |
|  | RIPAl5 | XM_019681673/**AALF015062** | KXJ73764 |
|  | RIPAl6 | **AALF016445** | KXJ73133 |
|  | RIPAl7 | **AALF016444** | KXJ73132 |
| *Bemisia tabaci* | RIPBt1 | XM_019046661 | XP_018902206 |
|  | RIPBt2 | XM_019045743 | XP_018902288 |
| *Bradysia odoriphaga* | RIPBo1 | from 40121 to 41143 of JAFDOW010000841 | KAG4072827 |
|  | RIPBo2 | from 6401 to 6401 of JAFDOW010000841 | Not annotated |
| *Contarinia nasturtii* | RIPCn1 | from 2078 to 3118 of VYII01002082 | Not annotated |
|  | RIPCn2 | XM_031770132 | XP_031625992 |

**Supplementary table 2.** Sequences Read Archives (SRA) used to analyze the expression of RIP genes across developmental stages of *A. aegypti*, *A. albopictus* and *C. nasturtii* are indicated in first, third and fifth columns, respectively. Evaluated stages of Embryos, Larval 1-4 (L1-L4), Early pupal (EP), Mid pupal (MP), Late pupal (LP) and Adult (A) are indicated in second, four and sixth columns. In some stages both sexes female (F) and male (M) were evaluated.

| **Developmental stages** | | | | | |
| --- | --- | --- | --- | --- | --- |
| *Aedes aegypti PRJNA419241* | | *Aedes albopictus PRJNA275727* | | *Contarinia nasturtii PRJNA565761* | |
| SRA files | Stage | SRA files | Stage | SRA files | Stage |
| SRX3411662, SRX3411661, SRX3411654, SRX3411653 | L1 | SRX885419 | Embryos | SRX6853820 | Embryos |
| SRX3411630, SRX3411629, SRX3411627 | L2 | SRX885422 | L1-L2 | SRX6853817 | L1 |
| SRX3411631, SRX3411626, SRX3411625, SRX3411623 | L3M | SRX885423 | L3-L4 | SRX6853818 | L2 |
| SRX3411632, SRX3411628, SRX3411624 | L3F | SRX885421 | P | SRX6853819 | L3 |
| SRX3411608SRX3411607, SRX3411606, SRX3411605 | L4M | SRX882870 | AM | SRX6853823 | P |
| SRX3411610, SRX3411609, SRX3411604, SRX3411603 | L4F | SRX885420 | AF | SRX6853822 | AM |
| SRX3411660, SRX3411659, SRX3411658, SRX3411657 | EPM |  |  | SRX6853821 | AF |
| SRX3411656, SRX3411655, SRX3411622, SRX3411621 | EPF |  |  |  |  |
| SRX3411640, SRX3411639,  SRX3411638, SRX3411637, SRX3411633 | MPM |  |  |  |  |
| SRX3411636, SRX3411635, SRX3411634 | MPF |  |  |  |  |
| SRX3411652, SRX3411651, SRX3411650, SRX3411646, SRX3411645 | LPM |  |  |  |  |
| SRX3411649, SRX3411612, SRX3411611 | LPF |  |  |  |  |
| SRX3411620, SRX3411619, SRX3411618, SRX3411617 | AM |  |  |  |  |
| SRX3411616, SRX3411615, SRX3411614, SRX3411613 | AF |  |  |  |  |

**Supplementary table 3.** SRA used to analyze the expression of RIP genes in different body parts of *A. aegypti*, and *B. tabaci* are indicated in first and third columns, respectively. Body parts analyzed are indicated in second and fourth columns.

| **Organism part** | | | |
| --- | --- | --- | --- |
| *Aedes aegypti* PRJNA236239 | Body part | *Bemisia tabaci* PRJEB26594 | Body part |
| SRX468784, SRX468785, SRX468786, SRX468787, SRX468788 | Female Antennae | ERX2611174 | Whole body |
| SRX468789, SRX468790, SRX468791, SRX468792 | Female Brain | ERX2611172, ERX2611173 | Thorax |
| SRX468793, SRX468794, SRX468795 | Female Rostrum | ERX2611176 | Abdomen |
| SRX468796, SRX468797, SRX468798 | Female Abdominal tip | ERX2611177 | Salivary glands |
| SRX468799, SRX468800, SRX468801, SRX468802, SRX468803, SRX468804 | Female Hindlegs | ERX2611178 | Midgut |
| SRX468805, SRX468806, SRX468807 | Female Midlegs |  |  |
| SRX468808, SRX468809, SRX468810 | Female Forelegs |  |  |
| SRX468719, SRX468720, SRX468721, SRX468722 | Female Maxilary palp |  |  |
| SRX468723, SRX468724, SRX468725, SRX468726, SRX468727 | Female proboscises |  |  |
| SRX468728, SRX468729, SRX468730, SRX468731, SRX468732, SRX468733 | Ovaries |  |  |
| SRX468761, SRX468762, SRX468763, SRX468764, SRX468765, SRX468766 | Male Antennae |  |  |
| SRX468755, SRX468756, SRX468757, SRX468758, SRX468759, SRX468760 | Male Brain |  |  |
| SRX468847, SRX468848, SRX468849, SRX468850, SRX468851, SRX468852 | Male Rostrum |  |  |
| SRX468776, SRX468777, SRX468778, SRX468779, SRX468780, SRX468781 | Male Abdominal tip |  |  |
| SRX468770, SRX468771, SRX468772, SRX468853, SRX468854 | Male Hindlegs |  |  |
| SRX468773, SRX468774, SRX468775 | Male Midlegs |  |  |
| SRX468767, SRX468768, SRX468769 | Male Forelegs |  |  |

**Supplementary table 4.** SRA used to analyze the expression of RIP genes present in *A. aegypti* after the infection with *B. malayi*. The conditions infected and uninfected for each strain are indicated in the second and fourth columns, respectively.

| ***Aedes aegypti* infected with *Brugiamalayi*** | | | |
| --- | --- | --- | --- |
| PRJNA255467 | Condition | PRJNA232599 | Condition |
| SRX665355, SRX665357, SRX667052 | 0 hs LVP-IB12^R^ | SRX399528 | BEY-LVP Day0 |
| SRX667053, SRX667054, SRX667055 | 0hs LVP-FR3^S^ | SRX399529 | BEY-LVP Day1 Uninf |
| SRX667056, SRX667057 | 12 hs infected LVP-IB12^R^ | SRX399530 | BEY-LVP Day2 Uninf |
| SRX667058, SRX667059, SRX667060 | 12 hs uninfected LVP-IB12^R^ | SRX399531 | BEY-LVP Day3 Uninf |
| SRX667061, SRX667062 | 12 Hs infected LVP-FR3^S^ | SRX399532 | BEY-LVP Day4 Uninf |
| SRX667063, SRX667064, SRX667065 | 12 Hs uninfected LVP-FR3^S^ | SRX399533 | BEY-LVP Day5 Uninf |
| SRX667066, SRX667067, SRX667068, SRX667069 | 48 hs infected LVP-IB12^R^ | SRX399534 | BEY-LVP Day6 Uninf |
| SRX667070, SRX667071, SRX667072, SRX667073 | 48 hs uninfected LVP-IB12^R^ | SRX399535 | BEY-LVP Day7 Uninf |
| SRX667074, SRX667075, SRX667076, SRX667077 | 48 Hs infected LVP-FR3^S^ | SRX399536 | BEY-LVP Day8 Uninf |
| SRX667078, SRX667079, SRX667080, SRX667081 | 48 Hs uninfected LVP-FR3^S^ | SRX399537 | RED Day0 Uninf |
|  |  | SRX399538 | RED Day1 Uninf |
|  |  | SRX399539 | RED Day2 Uninf |
|  |  | SRX399540 | RED Day3 Uninf |
|  |  | SRX399541 | RED Day4 Uninf |
|  |  | SRX399516, SRX399504 | BEY-LVP Day1 Inf |
|  |  | SRX399517, SRX399505 | BEY-LVP Day2 Inf |
|  |  | SRX399518, SRX399506 | BEY-LVP Day3 Inf |
|  |  | SRX399519, SRX399507 | BEY-LVP Day4 Inf |
|  |  | SRX399520, SRX399508 | BEY-LVP Day5 Inf |
|  |  | SRX399521, SRX399509 | BEY-LVP Day6 Inf |
|  |  | SRX399522, SRX399510 | BEY-LVP Day7 Inf |
|  |  | SRX399523, SRX399511 | BEY-LVP Day8 Inf |
|  |  | SRX399524, SRX399512 | RED Day1 Inf |
|  |  | SRX399525, SRX399513 | RED Day2 Inf |
|  |  | SRX399526, SRX399514 | RED Day3 Inf |
|  |  | SRX399527, SRX399515 | RED Day4 Inf |

1	Sikora, T., Jaschhof, M., Mantič, M., Kaspřák, D. & ševčík, J. Considerable congruence, enlightening conflict: molecular analysis largely supports morphology-based hypotheses on Cecidomyiidae (Diptera) phylogeny. *Zoological Journal of the Linnean Society* **185**, 98-110 (2019).
